# A mixture of sparse coding models explaining properties of face neurons related to holistic and parts-based processing

**DOI:** 10.1101/086637

**Authors:** Haruo Hosoya, Aapo Hyvärinen

## Abstract

Experimental studies have revealed evidence of both parts-based and holistic representations of objects and faces in the primate visual system. However, it is still a mystery how such seemingly contradictory types of processing can coexist within a single system. Here, we propose a novel theory called mixture of sparse coding models, inspired by the formation of category-specific subregions in the inferotemporal (IT) cortex. We developed a hierarchical network that constructed a mixture of two sparse coding submodels on top of a simple Gabor analysis. The submodels were each trained with face or non-face object images, which resulted in separate representations of facial parts and object parts. Importantly, evoked neural activities were modeled by Bayesian inference, which had a top-down explaining-away effect that enabled recognition of an individual part to depend strongly on the category of the whole input. We show that this explaining-away effect was indeed crucial for the units in the face submodel to exhibit significant selectivity to face images over object images in a similar way to actual face-selective neurons in the macaque IT cortex. Furthermore, the model explained, qualitatively and quantitatively, several tuning properties to facial features found in the middle patch of face processing in IT as documented by Freiwald, Tsao, and Livingstone (2009). These included, in particular, tuning to only a small number of facial features that were often related to geometrically large parts like face outline and hair, preference and anti-preference of extreme facial features (e.g., very large/small inter-eye distance), and reduction of the gain of feature tuning for partial face stimuli compared to whole face stimuli. Thus, we hypothesize that the coding principle of facial features in the middle patch of face processing in the macaque IT cortex may be closely related to mixture of sparse coding models.

## Introduction

The variety of objects that we see everyday is overwhelming and how our visual system deals with such complexity is a long-standing problem. Classical psychology has often debated on whether an object is represented as a combination of individual parts (parts-based processing) or as a whole (holistic processing) [1]. Experimental studies havexg348 cl:1 x revealed evidence of both types of processing in behaviors [1, 2] and in neural activities in higher visual areas [2–5], somewhat favoring holistic representation for faces and parts-based representation for non-face objects [1, 2, 5]. However, a theoretical question is: how could a single system reconcile such two seemingly contradictory types of processing? Although a number of studies on computational vision models showed remarkable performance in visual recognition [6–10], success in modeling higher visual areas [11, 12], or account for behavioral experiments on holistic face processing [12, 13], none of these studies offered insight into the tension between parts-based and holistic processing in a comparative manner with neurophyisology.

In this study, we address this question in a novel theoretical framework, called mixture of sparse coding models. We assume two separate sparse coding models, one dedicated to encode face images and the other to encode non-face object images, that perform competitive interaction. Sparse coding is well known for its close relationship with representations in early visual areas [14–22]; we transfer this technique to the study of higher visual representations. That is, exploiting the fact that sparse coding to image data of a specific category can yield parts-based feature representations (cf. [23, 24]), we constructed two separate category-specific representations for faces and objects analogously to the formation of specialized subregions for faces and objects in the inferotemporal (IT) cortex [25, 26]. Furthermore, we combined the two sparse coding models into a mixture model and modeled neural activities in terms of Bayesian inference. Then, we found that this framework gave rise to a form of holistic computation: not only recognition of the whole object depends on the individual parts, but also recognition of a part depends on the whole. This is in fact a Bayesian explaining-away effect: an input image is first independently interpreted by each sparse coding submodel, but then the one offering the better interpretation is adopted and the other is dismissed. For example, even if a part of an input image is a potential facial feature (e.g., a half-moon-like shape 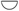), that feature would not be recognized as an actual facial feature (e.g., a mouth) if the whole image is a non-face object (Figure 1B).

**Figure 1.**
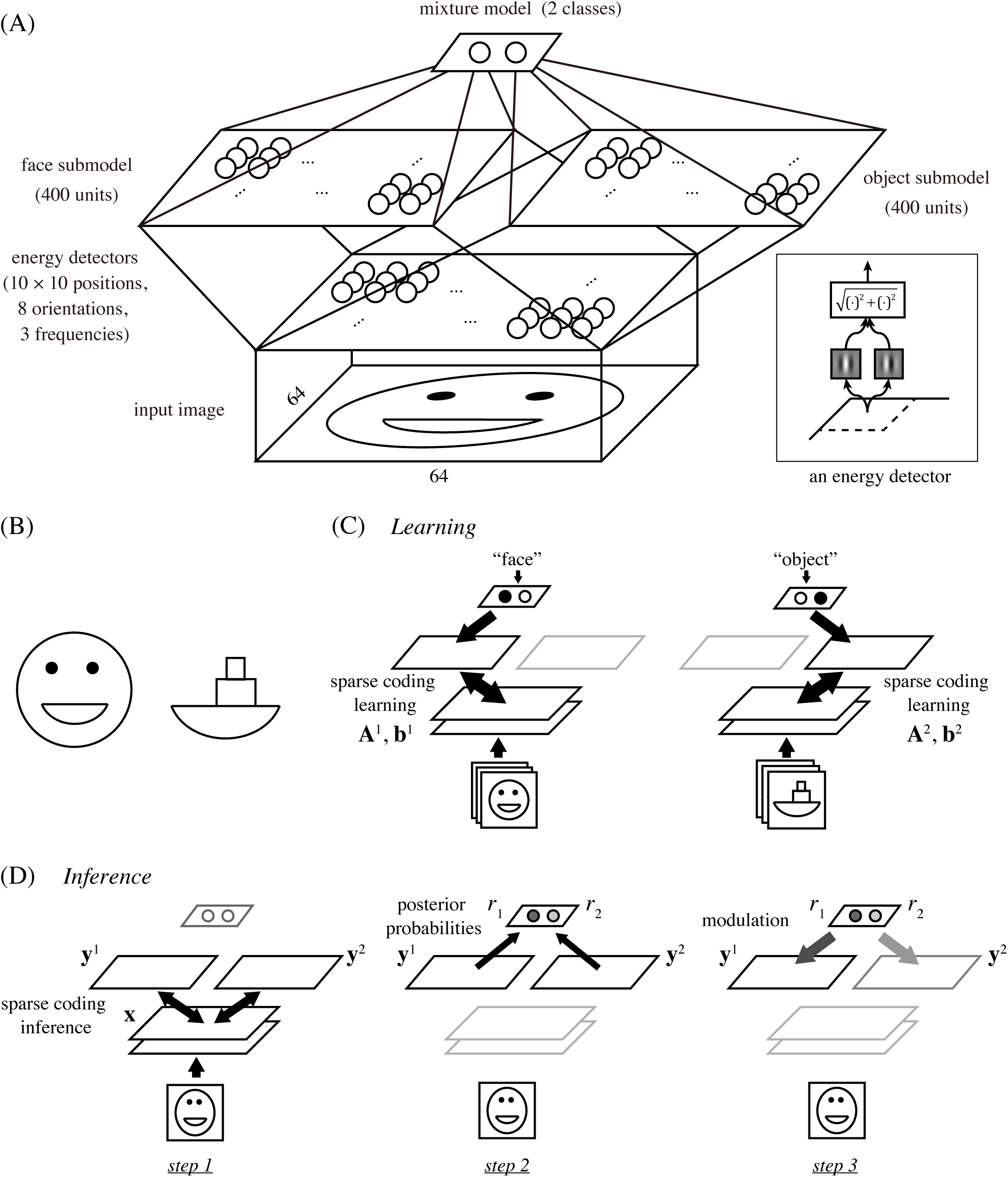
(A) The architecture of our hierarchical model. It starts with an energy detector bank and proceeds to two sparse coding submodels for faces and objects, which are then combined into a mixture model. Inset: an energy detector model. (B) Cartoon face and boat. Note that the mouth of the face and the base of the boat are the same shapes. (C) Learning scheme. We assume explicit class information, either “face” or “object,” of input images to be given during training, which allows us to use a standard sparse coding learning for each submodel with the corresponding dataset. (D) Inference scheme. For testing response properties, the network first interprets the input separately by the sparse code of each submodel (step 1), then compares the goodnesses of the obtained interpretations as posterior probabilities (step 2), and finally modules multiplicatively the responses in each submodel with the corresponding posterior probability (step 3).

We discovered that our model had a close relationship with computation known for a region of the macaque IT cortex called the face-selective middle patch, as documented by Freiwald et al. [4]. First, our model cells in the face submodel exhibited prominent selectivity to face images over non-face object images in a similar way to actual face-selective neurons, and this selectivity was crucially dependent on the above-mentioned explaining-away effect. Second, these model cells reproduced a number of tuning properties of face neurons in the middle patch. In particular, our model face cells tended to (a) be tuned to only a small number of facial features, often related to geometrically large parts such as face outline and hair, (b) prefer one extreme for a particular facial feature while anti-prefering the other extreme, and (c) reduce the gain of tuning when a partial face was presented compared to a whole face. We quantified these properties and compared these with the experimental data at the population level [4]; the result showed a good match. Thus, we propose the hypothesis that regions of the IT cortex representing objects or faces may employ a computational principle similar to mixture of sparse coding models.

## Results

### Model

To investigate the computational principles underlying face and object processing in the IT cortex, we designed a multi-layer network model illustrated in Figure 1A. The network had the architecture that received an image of 64 × 64 pixels, processed it with a fixed bank of standard energy detector models, and fed the results to two sparse coding models, called face submodel and object submodel (each with 400 model neurons), which were then combined into a mixture model to perform competitive interaction as explained later.

Each energy detector computed the squared norm of the outputs from two Gabor filters for the input image (Figure 1A, inset). The two filters had the same center position, orientation, and spatial frequency, but had phases different by 90°. The entire bank of energy detectors had all combinations of 10 × 10 center positions (in a grid layout), 8 orientations, and 3 frequencies; thus, the output of this stage had a total of 2400 dimensions (see the section on Model details in Methods.) In the actual visual cortex, inputs to IT areas are presumably computed between V1 and V4 and this computation must be much more complex than the energy detector bank in our model. However, some important aspects should still be reflected by this simple operation since a large number of V4 neurons are known to be orientation-selective [27]; moreover, this simple assumption was sufficient to reproduce certain response properties of face neurons as shown in what follows.

In training the mixture model, we assumed, for simplicity, that the class label of each input image, either “face” or “object,” was given (Figure 1C). This allowed us to use a naive learning procedure that separately trained each face or object submodel with an existing sparse coding method. Specifically, we used publicly available face and object image datasets in which the faces or objects were properly aligned within each image frame [24, 28, 29] (see the section on Data preprocessing in Methods). Then, for each image class *k*, which was either 1 (face) or 2 (object), we learned the basis matrix **A***^k^* and the mean vector **b***^k^* by sparse coding of the corresponding set of images that were processed by the energy model. (The basis matrix and mean vector were used for determining the responses of the model neurons to an input as explained below.) Classical mixture models are usually trained with an unsupervised learning method without class labels [30]. However, such learning is generally not easy and not our main interest here since we focus on inference, i.e., on computation of evoked responses, not on learning or plasticity. (We come back to this point in the Discussion section.)

To perform sparse coding learning, we adopted our previously developed approach based on independent component analysis (ICA) [22], which is known to be a good approximation of sparse coding [31] and for which efficient algorithms exist. In this approach, an important step was to drastically reduce the input dimensions, from 2400 to 100 dimensions here, by principal component analysis (PCA) before performing ICA. This is, in fact, a simple modification of a standard preprocessing used in any classical sparse coding or ICA methods. However, we have previously discovered that such strong dimension reduction has an effect of spatial pooling [32] and thereby produces much larger basis patterns than without it [22]. In the present case, we later show that weaker dimension reduction resulted in representations of overly small features, which led to a loss of discriminative power. After this step, to regain enough components from the reduced dimensions, we used overcomplete ICA [33], estimating 400 components from 100 dimensions. (See the section on Learning details in Methods.)

Once the network was trained, the response properties of the model neurons were tested using various input images. In this phase, we never explicitly gave class information on each input image, but rather let the network estimate it by Bayesian inference, which worked in the following three steps (Figure 1D).

1. Given an input **x** (processed by the energy detectors), interpret it separately by each submodel *k*. Formally, infer the responses 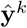 in each submodel *k* that maximize the sparse coding objective *L_k_*:

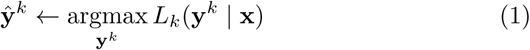

where

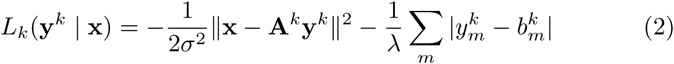

using pre-fixed constants σ and λ. Recall that **A***^k^* and **b***^k^* are the basis matrix and the mean vector for submodel *k* that are obtained in the learning phase as described above.
2. Compare the goodnesses of the two interpretations in the form of posterior probabilities. Formally, for each *k*:

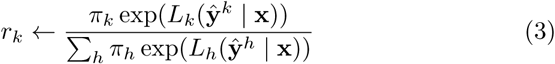

using a pre-fixed constant π*_k_* for prior probability. For simplicity, we assume π*_k_* = 1/2.
3. Modulate multiplicatively the responses in each submodel by the corresponding posterior probability computed above. That is, for each *k*:

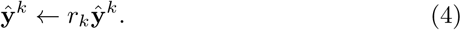

Step 1 is similar to inference in the classical sparse coding [31], where the responses in each submodel are estimated so as to minimize the reconstruction error and maximize the sparsity at the same time. One difference is, however, that the sparsity constraint here is on the difference from the mean vector **b***^k^*. We assume here a non-zero mean since the mean of face images is not zero and such stimulus usually elicits non-zero responses of actual face neurons, while the classical sparse coding assumes a zero mean since the mean of natural image patches is a blank, gray image, and such stimulus evokes no response of V1 neurons. The last two steps in our inference are a major departure from the classical sparse coding, where step 2 computes the posterior probability indicating how well each submodel interprets the input and step 3 multiplies the responses in each submodel by the corresponding posterior probability. By these steps, even if the input contains a feature that can potentially activate some units in a submodel, such units may eventually be deactivated when the whole input was not interpreted well by this submodel compared to the other submodels (Bayesian explaining-away effect).

Finally, to compare with neural responses later, we passed the response value of each unit (after step 3) to the smooth half-wave rectifying function *h*(*a*) = log(1 + exp(*a*)), which always produces non-negative values.

Although we presented above the mixture model and its inference computation in an informal and procedural way, these can be formalized rigorously within a probabilistic generative model. Generally, the motivation for such formalization is to regard visual recognition as a process of inferring hidden causes in the external world that generate a natural image. Our model can be seen as one such approach: all the computations described above can be derived from Bayesian inference of posterior probabilities in a statistical framework of mixture of sparse coding models. The details can be found in the section on Theory of mixture of sparse coding models in Methods.

### Basis representations

We proceed to show the representation in our model obtained by the learning procedure described so far. The basis matrix **A***^k^* of each submodel defines its internal representation and each column vector of the matrix (basis vector) exposes the specific feature represented by each unit. Figure 2 shows the basis vectors of three example units in the face submodel. Each unit is visualized as a set of ellipses corresponding to the energy detectors, where their underlying Gabor filters have the indicated center positions (in the visual field coordinates), orientations, and spatial frequencies (inversely proportional to the size of the ellipse). The color of the ellipse indicates the weight value normalized by the maximal weight value. For readability, we show only the ellipses corresponding to the maximal positive (excitatory) weight and the minimal negative (inhibitory) weight at each location. Although this visualization approach may seem a bit too radical, it did not lose much information: we confirmed by visual inspection that the local weight patterns for most units had only one positive peak and one negative peak at each position and frequency and the patterns of orientation integration did not have notable changes across frequencies. In Figure 2, we can see that unit #1 represented a face outline either on the left (excitatory) or on the right (inhibitory); unit #2 represented mainly eyes (excitatory); unit #3 mainly represented a mouth (excitatory) and nose (weakly inhibitory). Figure 3 shows the basis vectors of 32 randomly selected units from (A) the face submodel and (B) the object submodel. The representations in these two submodels were qualitatively different: face units represented local facial features (i.e., facial parts like outline, eye, nose, and mouth) and object units represented local object features.

**Figure 2.**
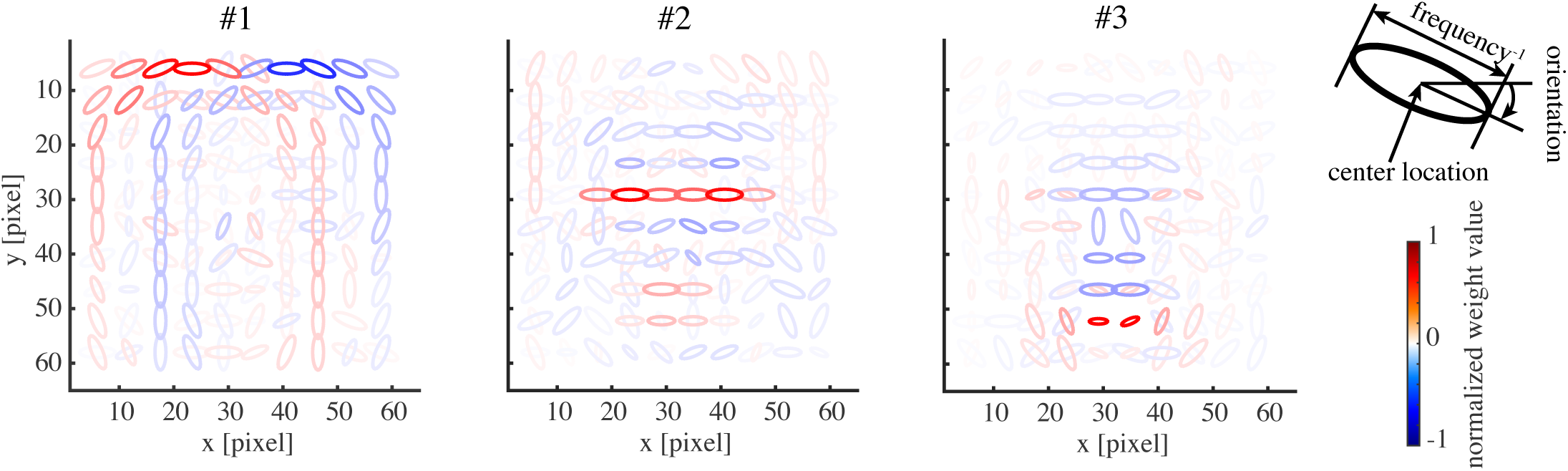
The basis representations of three sample model face units. Each panel depicts the weighting pattern (basis vector) from a face unit to energy detectors by a set of ellipses, where each ellipse corresponds to the energy detector at the indicated x-y position, orientation, and frequency (inverse of the ellipse size); see the top right legend. The color shows the normalized weight value (color bar). Only the maximum positive and the minimum negative weights are shown at each position for readability.

**Figure 3.**
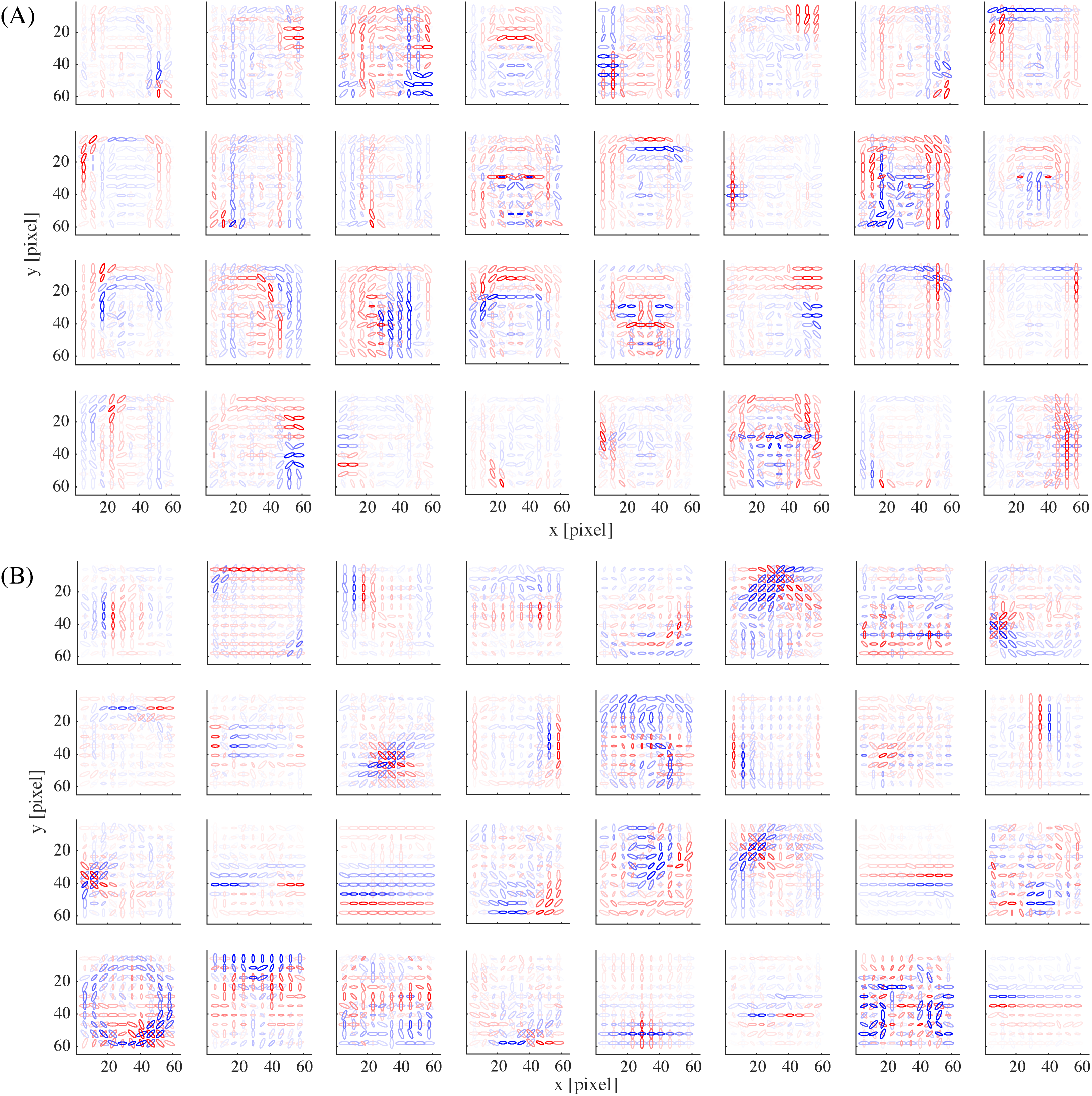
The basis representations of (A) 32 example model face units and (B) 32 example model object units.

### Selectivity to faces

Next, we show a series of comparisons between the response properties of our model and the experiments conducted by Freiwald et al. [4] on the region in monkey IT cortex called the face middle patch.

As mentioned above, due to the Bayesian explaining-away effect in the mixture model, model face units exhibited selectivity to face images and object units to object images. We measured the responses of our model units to natural face and object images that were separate from the training images (without explicitly giving class labels). The left panel of Figure 4A shows the responses (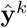 in step 3 of Bayesian inference) of the face units (top) and object units (bottom) to face images, where the images were sorted by the response magnitudes, separately for each unit. The right panel similarly shows the responses of the same units to object images. We can see that the face units were prominently responsive to many face images while indifferent to non-face object images; the object units had the opposite property. To quantify such face selectivity, we calculated the face-selectivity index for each unit, which was defined as the ratio between the difference and the sum of the mean response to faces and the mean response to objects (where the baseline, i.e., the response to a blank image, was subtracted from each response value). Figure 4D (blue) shows the distribution of face-selectivity indices for the face units. The result indicates almost no unit with index between −1/3 and −1/3, which is consistent with the experimental data [4, Figure 1b].

**Figure 4.**
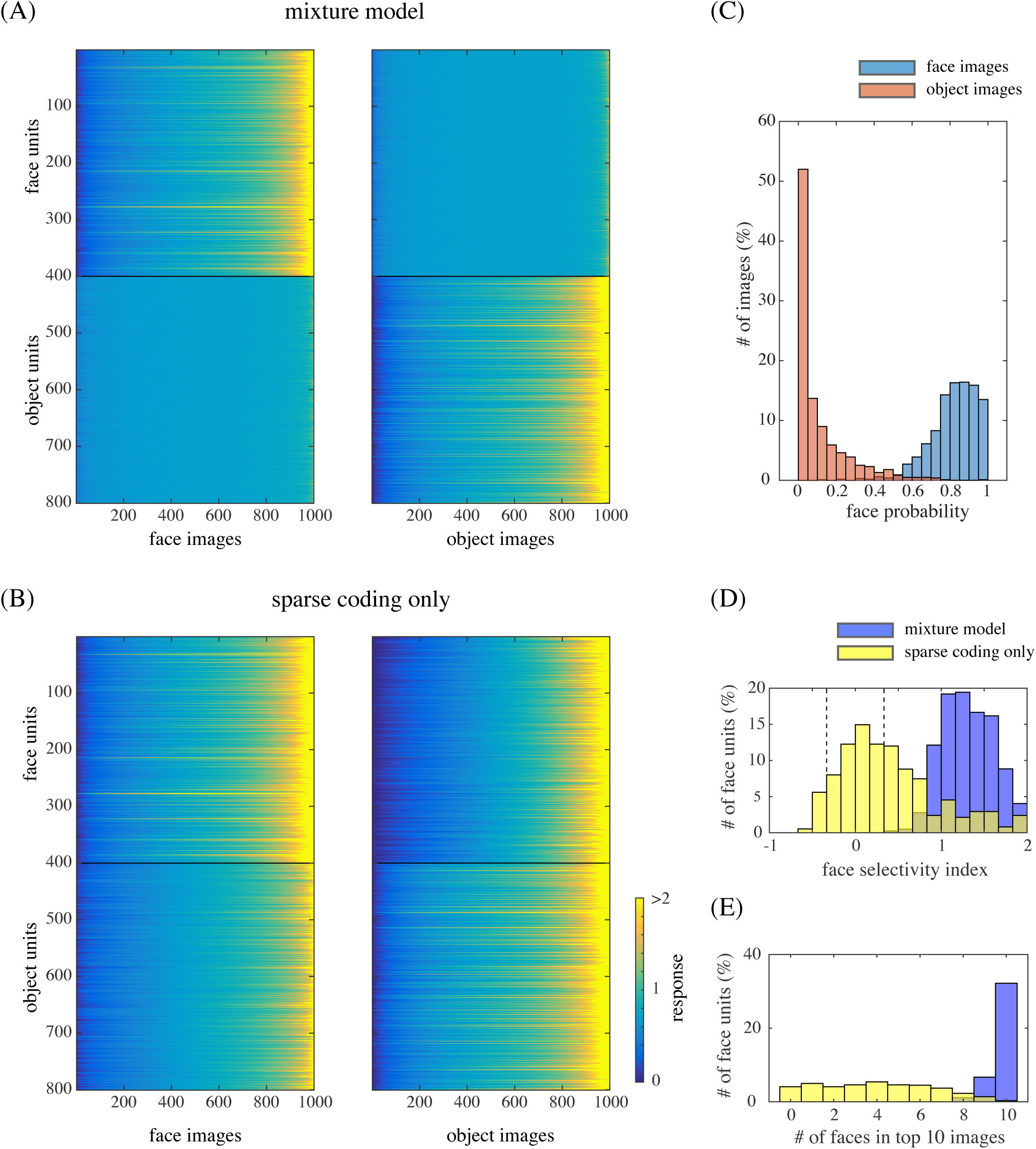
(A) The responses of model face units (1–400) and model object units (401–800) to face images (left) and object images (right). The images are sorted by response magnitudes (color bar) for each unit. (B) The responses in the case of removing mixture computation. (C) The distribution of face posterior probabilities for face image inputs and for object image inputs. (D) The distribution of face-selectivity indices for the face units in the case of the mixture model (blue) or the case of the sparse coding model (yellow). The broken lines indicate the values −1/3 and 1/3. (E) The distribution of the number of face images in the top 10 (face or object) images that elicited the largest responses of each face unit.

Such vivid selectivities disappeared when the mixture computation was removed. Figure 4B shows the analogous responses of the face and object units immediately after performing sparse coding (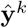 in step 1); the face units became almost equally responsive to object images to face images. Indeed, Figure 4D (yellow) shows that the face-selectivity indices of those units became substantially lower by the removal of mixture, with a majority falling between −1/3 and 1/3.

To gain more insight into the underlying computations, see the distributions of face posterior probabilities (*r*_1_ in step 2) for face and object images in Figure 4C: faces and objects were clearly discriminated. In fact, those posterior probabilities modulated the response of each unit representing a part (step 3), which resulted in prominent face selectivity. (Note that the discrimination capability did not automatically arise from step 3 since it actually depended on proper training of both submodels; see the section on “Control simulations.”) Further, Figure 4E shows that the images that elicited the largest responses of the face units were mostly faces in the mixture model (blue), whereas it was not the case in the model without mixture (yellow). Thus, even though the face units by themselves could detect accidental features similar to facial parts, the mixture computation ensured that they responded only when the whole input was a face image. In other words, face selectivity can be interpreted as a form of holistic processing in our mixture model.

### Tuning to facial features

We next turn our attention to tuning properties to facial features. The experiment by Freiwald et al. [4] used cartoon face stimuli for which facial features were controlled by 19 feature parameters, each ranging from −5 to +5. The authors recorded responses of a neuron in the face middle patch while presenting a number of cartoon face stimuli whose feature parameters were randomly varied. Then, for each feature parameter, they estimated a tuning curve by taking the average of the responses to the stimuli that had a particular value while varying other parameters (“full variation”). We simulated the same experiment and analysis on our model (see the section on Simulation details in Methods; see also Figure S3 for examples of cartoon face images.).

To illustrate tuning to facial features in our model, Figure 5 shows the tuning curves of the face units in Figure 2 to all 19 feature parameters. Each unit was significantly tuned to one to nine feature parameters (where significance was defined in terms of surrogate data; see Methods). Some tunings clearly reflected the corresponding parts in the basis representations. Unit#1 was tuned only to the face direction, preferring the left as opposed to the right. Unit#2 mainly showed tuning to eye-related features, in particular, preferring narrower inter-eye distances and larger irises. Unit#3 mainly showed tuning to mouth-and nose-related features, in particular, preferring smily mouths and longer noses.

**Figure 5.**
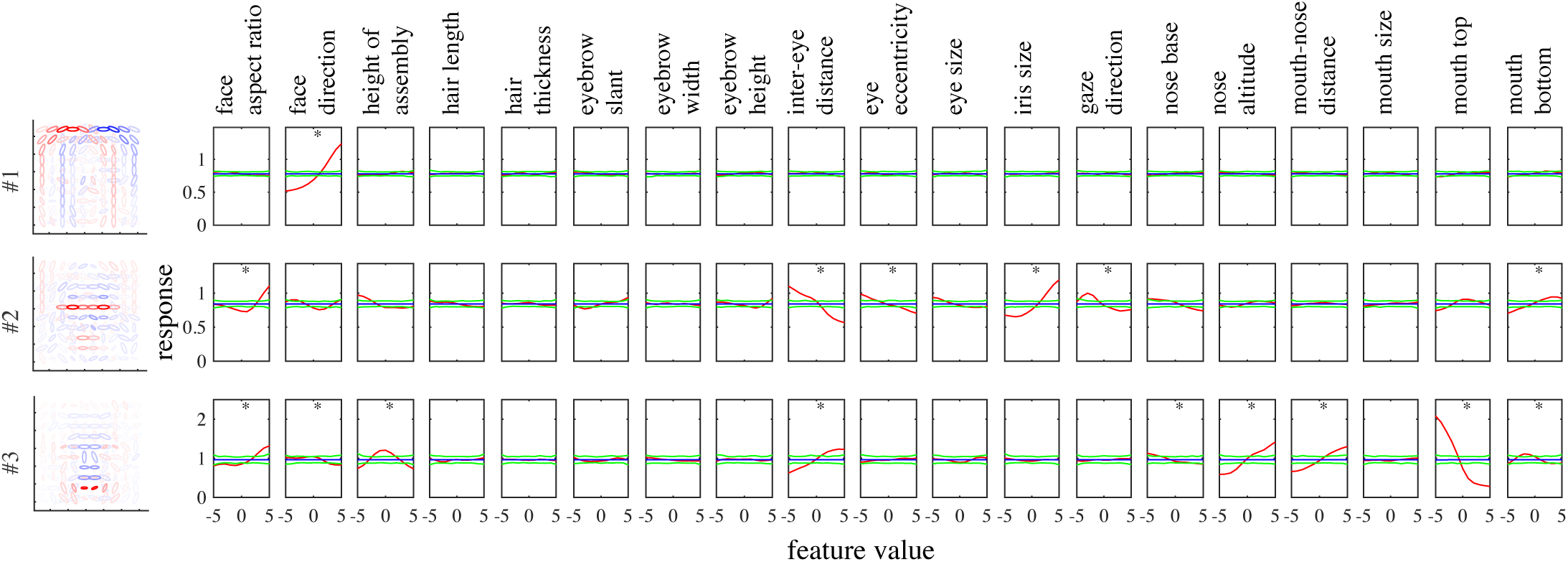
The tuning curves (red) of the model face units shown in Figure 2 to 19 feature parameters of cartoon faces. The mean (blue) as well as the maximum and minimum (green) of the tuning curves estimated from surrogate data are also shown (see the section on Simulation details in Methods).

Even in the whole population, most units were significantly tuned to only a small number of features similarly to the experiment [4]. Figure 6A shows the distribution of the numbers of tuned features per unit, which were on average 3.6 and substantially smaller than 19, the total number of features. The face neurons in the monkey face middle patch were also tuned to only a small number of features, i.e., 2.6 on average [4, Figure 3c] (replotted in red boxes in Figure 6A). Figure 6B shows the distribution of the numbers of significantly tuned units per feature. The distribution strongly emphasizes geometrically large parts, i.e., face aspect ratio, face direction, feature assembly height, and inter-eye distance. The shape of the distribution has a good match with the experimental result [4, Figure 3d] (replotted in Figure 6B), though iris size seems much more represented in the monkey case.

**Figure 6.**
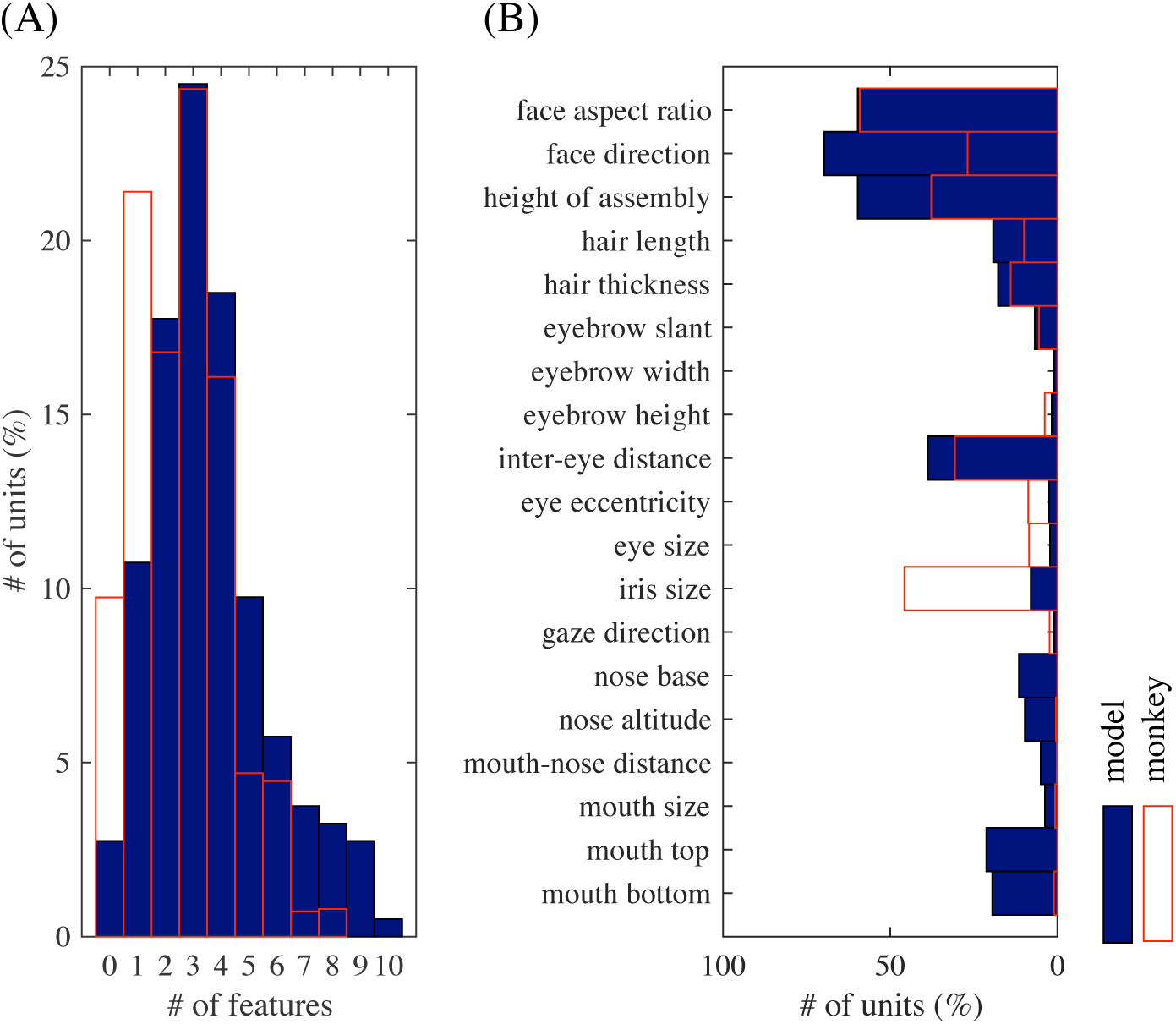
(A) The distribution of the numbers of significantly tuned features per unit, overlaid with a replot of [4, Fig. 3c]. (B) The distribution of the numbers of significantly tuned units for each feature parameter, overlaid with a replot (red boxes) of [4, Fig. 3d].

A prominent property of the experimentally obtained tuning curves was preference or anti-preference of extreme facial features [4]; our model reproduced this property as well. For example, Figure 5 shows that many tuning curves were maximum or minimum at one of the extreme values (−5 or +5). For the entire population, Figure 7A shows all significant tuning curves of all face units, sorted by the peak feature values. To quantify this, Figure 7B shows the distributions of peak and trough feature values; the extremity preference index (the ratio of the average number of peaks in the extreme values to the number of peaks in the non-extreme values) was 9.1 and the extremity anti-preference index (analogously defined for troughs) was 12.0. These indicate that the tendency of preference or anti-preference of extreme features generally held for the population. This result is in good agreement with the monkey experiment [4], which also reported distributions of peak and trough values that were biased to the extreme values [4, Fig. 4a] (the extremity preference indices were 7.0, 5.5, and 7.1, and the extremity anti-preference indices were 12.6, 13.7, and 12.1 for three monkeys; the average distribution is replotted in Figure 7B).

**Figure 7.**
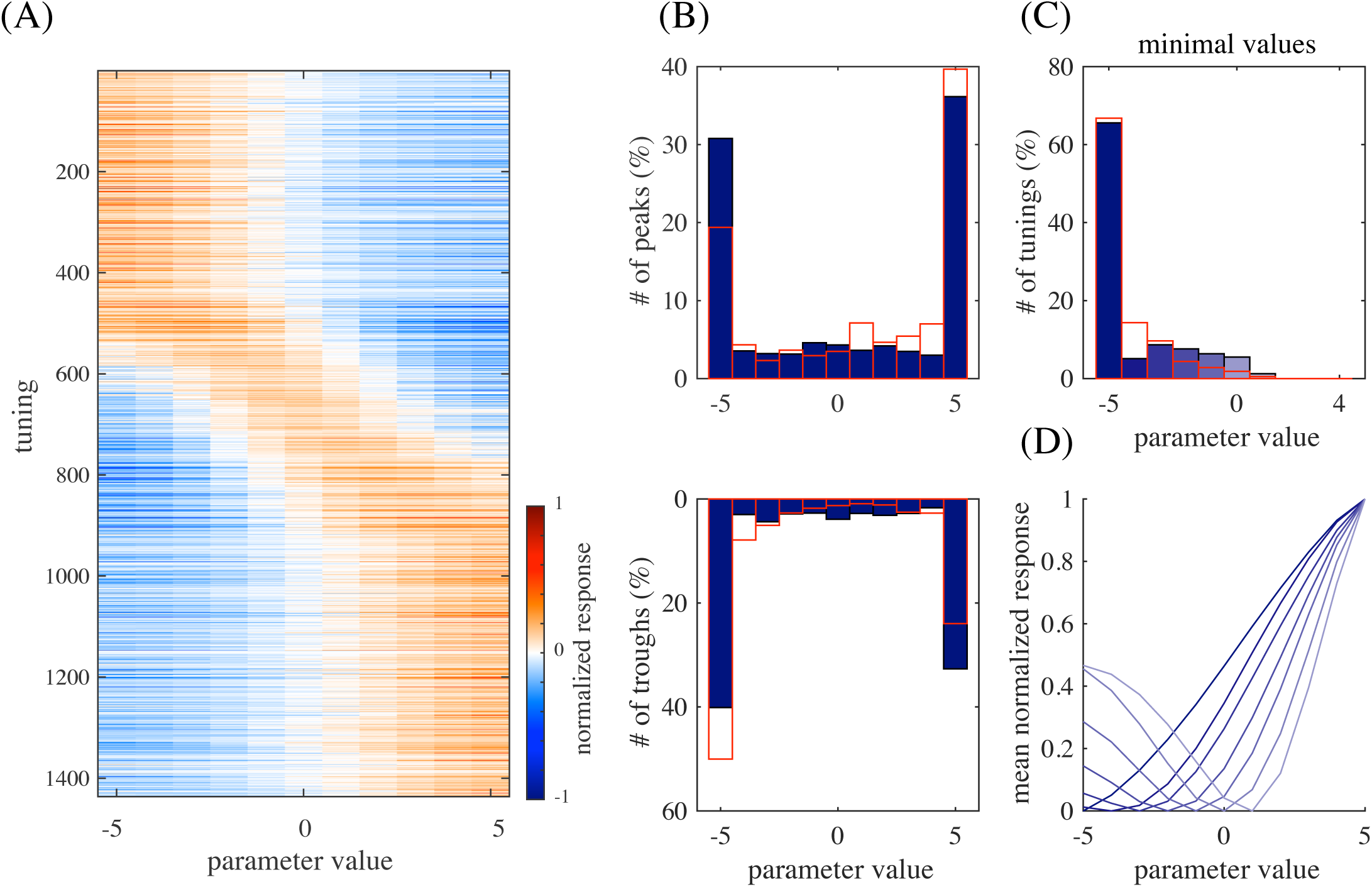
(A) All significant tuning curves of all model face units sorted by the peak parameter value. Each tuning curve (row) here was mean-subtracted and divided by the maximum. (B) The distributions of peak parameter values (top) and of trough parameter values (bottom). The overlaid red boxes are replots of [4, Fig. 4a] averaged over three monkeys. (C) The distribution of minimal values of the significant tuning curves peaked at +5 and the flipped tuning curves peaked at −5, overlaid with a averaged replot of [4, Fig. 4d]. (D) The average of the tuning curves for each minimal value in (C) (with the same color).

In addition, the experimental study even observed monotonic tuning curves [4], which were also found in our model as in Figure 5. To quantify this for the population, Figure 7C shows the distribution of minimal values of the significant tuning curves preferring value +5 pooled together with the tuning curves preferring value −5 that have then been flipped; the distribution has a clear peak at value −5. Further, for each minimal value in Figure 7C, the average of the tuning curves (normalized by the maximum response) with that minimal value is given in Figure 7D; the averaged tuning curve for minimal value −5 has a monotonic shape. These indicate that tuning curves preferring one extreme value tended to anti-prefer the other extreme value and be monotonic. This result is consistent with the experimental data, which also showed a distribution of minimal values that was peaked at −5 [4, Fig. 4d] (replotted in Figure 7C) and a monotonic averaged tuning curve corresponding to minimal value −5 [4, Fig. 4d, inset]. We discuss later why the model face units acquired such extremity preferences.

We have explained above the face selectivity property as a form of holistic processing in the mixture model. On the other hand, the experimental study investigated holistic face processing in the IT cortex by using partial face stimuli and inverted face stimuli [4]. To gain insight into these experiments, we also conducted simulations of the same experiments in our model.

To simulate the experiment with partial faces [4], we estimated two kinds of tuning curves in addition to the one used so far (“full variation”), namely, the responses to full cartoon faces where one feature was varied and the other were fixed to standard ones (“single variation”) and the responses to partial faces where only one feature was presented and varied (“partial face”). (See the section on Simulation details in Methods.) Figure 8 compares tuning curves in (A) full variation vs. single variation, (B) full variation vs. partial face, and (C) single variation vs. partial face. Overall, the shapes of the tunings were similar for all three kinds (average correlation 0.94 to 0.95). However, the gain of each tuning function (the slope of the fitted linear function) tended to drop after the removal of most of facial features (Figure 8C); the average gain ratio was 2.0, which was close to 2.2, the experimentally reported number [4, Fig. 6c]. This effect was not only because typical face units represented a combination of two features or more, but also because partial faces looked less face-like than full faces: Figure 8E shows lower face posterior probabilities for the partial face condition than the full variation condition. Indeed, such drop was weakened when the mixture computation was removed: the average gain ratio was 1.5 when the same comparison was made for the responses of model face units without the mixture computation, i.e., using only step 1 in Bayesian inference (Figure 8D). In addition to these, note that the tunings curves in full variation were slightly reduced compared to those in single variation (Figure 8A–B); a similar tendency can be observed in the experimental result [4, Fig. 6c]. This reduction in the model was because the face images used in the single variation condition took standard feature values for most parameters and such face images looked more face-like than others (giving slightly larger face posterior probabilities than the full variation condition; Figure 8E).

**Figure 8.**
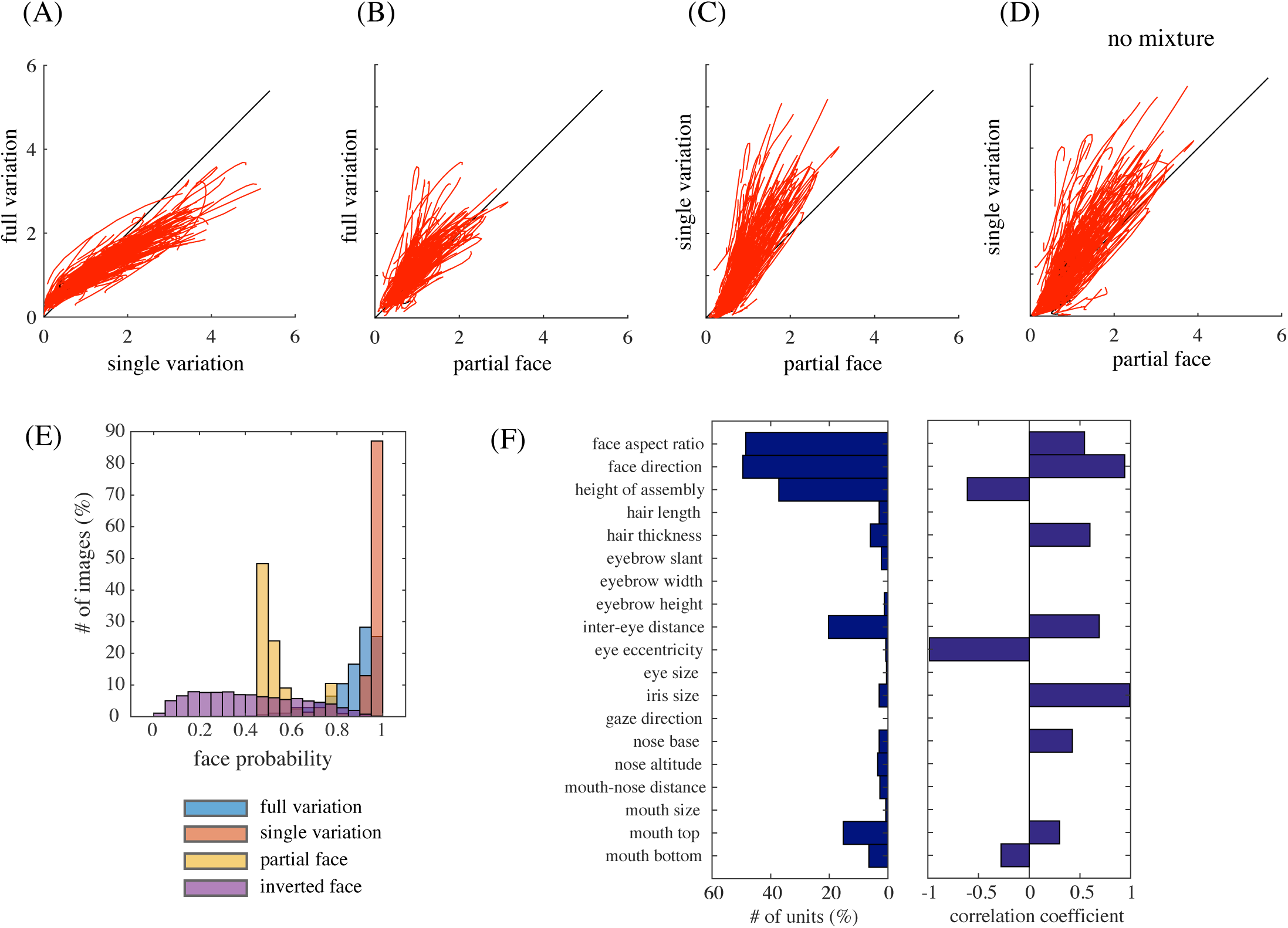
(A) Full-variation versus single-variation tuning curves. (B) Full-variation versus partial face tuning curves. (C) Single-variation versus partial face tuning curves. (D) Single-variation versus partial face tuning curves in the case of removing mixture computation. (E) The distributions of face posterior probabilities for the full variation, the single variation, the partial face, and the inverted face conditions. (F) The distribution of the numbers of tuned units per feature for inverted faces (left) and the mean correlation coefficient between the tunings for upright faces and for inverted faces for each facial feature (right).

To simulate the experiment with inverted faces [4], we presented, to the model, the same set of full cartoon faces except for their vertical inversion and estimated tuning curves for each facial feature in the same way (full variation). As a result, we found that the number of units that were tuned to each facial feature was more or less similar to the original model (Figure 8F, left). However, the tuning curves for assembly height tended to be inverted, whereas those for most other features did not (Figure 8F, right; for eye eccentricity, only two units had significant tunings and they happened to have a highly negative correlation between the upright and inverted cases). These results were consistent with the experiment [4, Figure 7ad]. However, we also observed that the overall responses of the model face units to inverted faces were much lower compared to upright faces (a somewhat similar tendency can be discernible in the experimental report [4, Figure 7bc]). This was because the mixture model could not classify well the inverted faces as faces since the face submodel was trained only with upright face images; consequently, the face posterior probabilities were generally low for inverted faces (Figure 8E, violet). Taken together, our result indicates that feature tuning for inverted faces could be explained by representation of individual parts of upright faces, although whole inverted faces may not be recognized as faces.

Interaction between feature parameters was limited, though present. For each pair of feature parameters, a 2D tuning was estimated by averaging the responses to a pair of parameter values while varying the remaining parameters. Then, the 2D tuning for a pair of parameters was compared to another 2D tuning predicted by the sum of two (full-variation) 1D tunings for the same parameters or by the product of these. The distributions of correlation coefficients are given in Figure 9; the averages were both 0.90, which was similar to the experimental result (averages 0.88 and 0.89) [4, Figure 5b].

**Figure 9.**
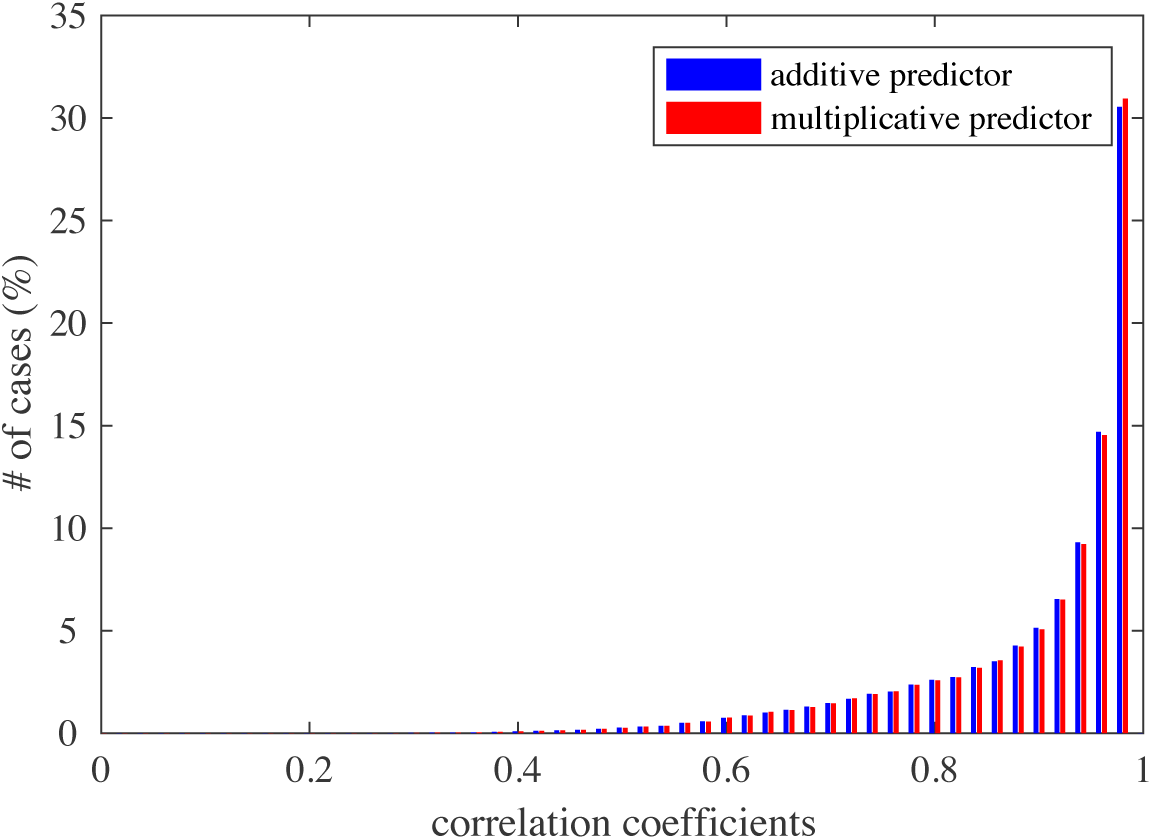
The distributions of correlation coefficients between 2D tuning functions and additive (blue) or multiplicative predictors (red).

### Control simulations

How much do our results depend on the exact form of model? To address this question, we modified the original model in various ways and conducted the same analysis.

First, we already showed that, when we omitted the mixture computation and simply used a sparse coding model of face images, the model units were deprived of selectivities to faces vs. objects (Figure 4). However, tuning properties to facial features did not change much. Figure 10 shows that the distributions of the number of tuned features per unit, of the number of tuned units per feature, of the peak feature values, and of the trough feature values for the modified model (cyan curves) are all similar to the original model (blue curves). Therefore, while the selectivities were from the mixture model, the tuning properties were produced by the sparse coding.

**Figure 10.**
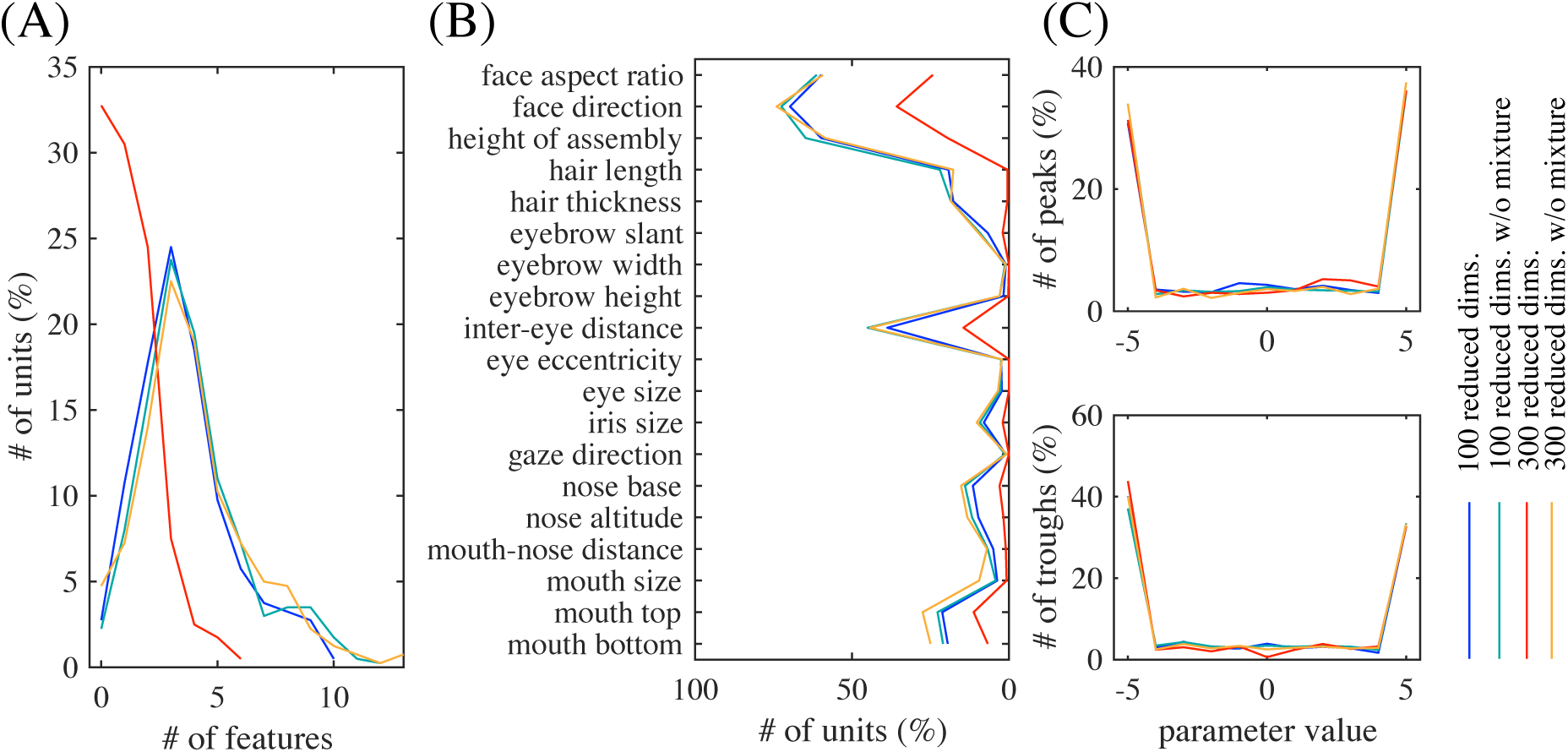
The distributions of (A) the number of tuned features per unit (cf. Figure 6A), (B) the number of tuned units per feature (cf. Figure 6B), and (C) the peak (top) and the trough (bottom) feature values (cf. Figure 7B), in different model variations. The color of each curve indicates the model variation (see legend).

Next, we varied the strength of dimension reduction of the outputs of the energy detector bank before performing sparse coding learning (the original model reduced the dimensionality from 2400 to 100). Three observations were made. First, consistently with our previous observation in our V2 model [22, 32], overall feature sizes tended to decrease while the reduced dimensionality was increased. Figure 12 shows example face and object units in the case of 300 reduced dimensions; compare these with Figure 3. (When we further increased the reduced dimensionality, we obtained quite a few units with globally shaped, somewhat noisy basis representations. These seemed to be a kind of “junk units” that are commonly produced when the amount of data is insufficient compared to the input dimensionality.) Second, as the reduced dimensionality increased, face posterior probabilities (as in Figure 4C) were substantially decreased for face images (Figure 11); the face images could barely be discriminated in the case of 300 reduced dimensions. Meanwhile, face posterior probabilities remained low for object images. This seemed to happen because the object submodel now learned to represent spatially very small and generic features so that it could give sufficiently good interpretations not only to object images but also to face images. This justified our model construction approach that performs strong dimension reduction before sparse coding learning. Third, Figure 10A–B shows that the number of tuned features per unit and the number of tuned units per feature decreased in the case of 300 reduced dimensions (red curve). This was due to the weakened selectivity rather than the size decrease of feature representations since the effect disappeared when the mixture computation was omitted (yellow curve).

**Figure 11.**
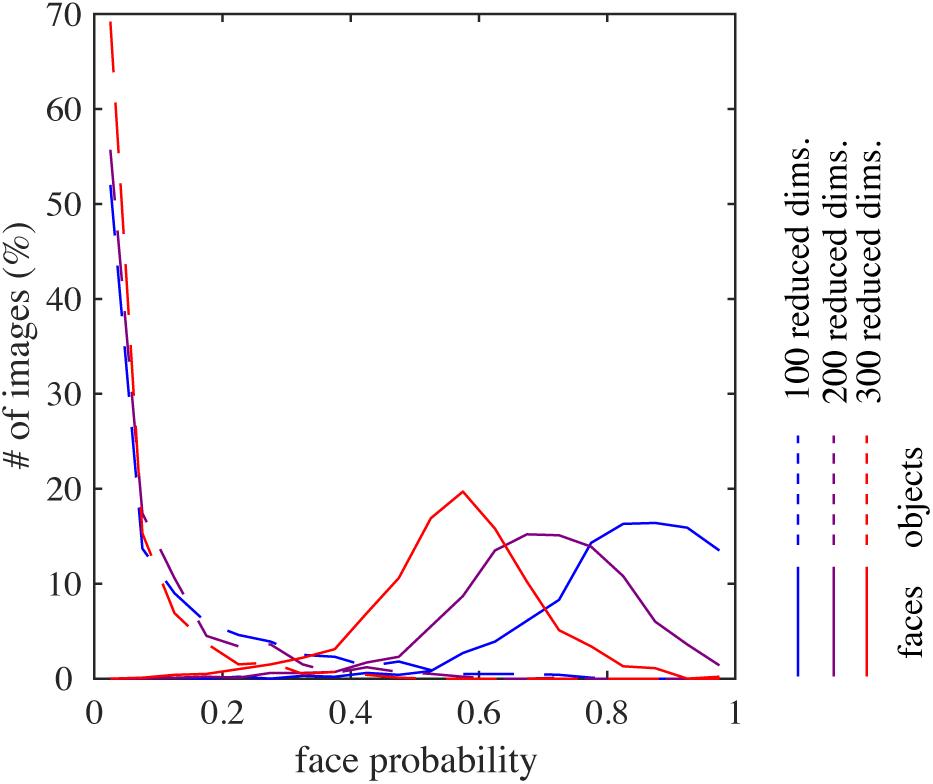
The distribution of face posterior probabilities for face images (solid curve) or for object images (broken curve) in different model variations (cf. Figure 4C). The color of each curve indicates the model variation (see legend).

**Figure 12.**
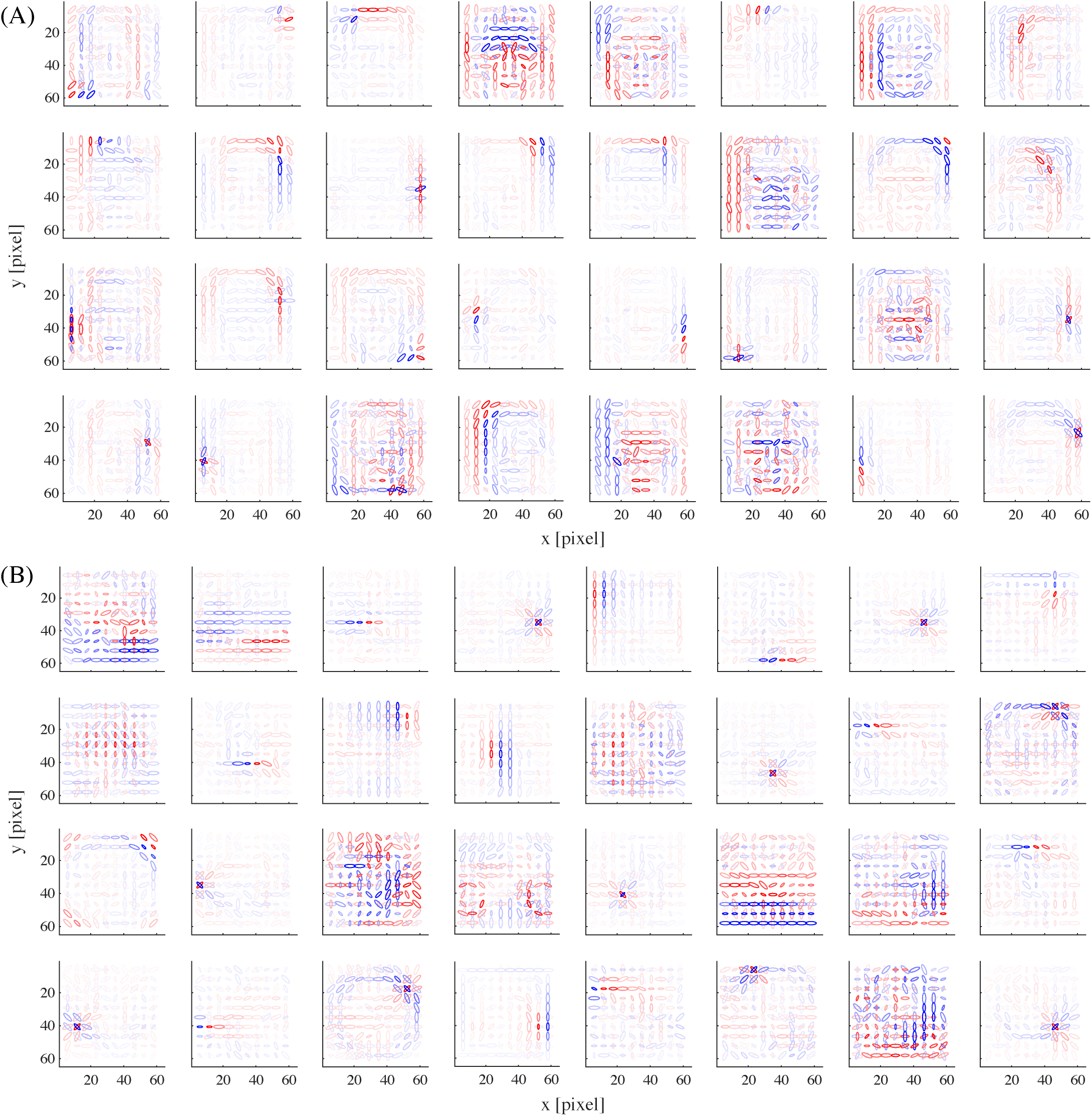
The basis representations of 32 example model units from (A) the face submodel and (B) the object submodel, in the network trained with 300 reduced dimensions.

As an additional control simulation, we varied the number of units (200 or 800) in each submodel of the mixture model while keeping the other conditions. In either case, we observed no discernible difference in the results from the original model (Figure S1).

We also examined a single sparse coding model (with no mixture model) trained with face and non-face images all together. In this model, we found almost no unit having face selectivity that was as vivid as in the original model; even for the units that gave average responses larger to faces than non-faces (which were only less than 10% of the whole population), selectivity to face images was rather weak, with face-selective indices mostly less than 1/3 (Figure S2A). However, such weakly face-selective units showed tuning properties similar to the original model (Figure S2B). Taken together, the response properties of those units were comparable to the sparse coding model trained only with faces without mixture model (Figure 4B and Figure 10, cyan curves).

## Discussion

In this study, we proposed a novel framework called mixture of sparse coding models and used this to investigate the computational principles underlying face and object processing in the IT cortex. In this model, two sparse feature representations, each specialized to faces or non-face objects, were built on top of an energy detector bank and combined into a mixture model (Figure 1). Evoked responses of units were modeled by a form of Bayesian inference, in which each sparse coding submodel attempts to interpret a given input by its code set, but the best interpretation explains away the input, dismissing the explanation offered by the other submodel. The model units in our face submodel not only exhibited significant selectivity to face images similarly to actual face neurons (Figure 4), but also reproduced qualitatively and quantitatively tuning properties of face neurons to facial features (Figures 5 to 9) as reported for the face middle patch, a particular subregion in the macaque IT cortex [4]. Thus, computation in this cortical region might be somehow related to mixture of sparse coding models.

While sparse coding produced parts-based representations in each submodel (Figures 2 and 3), the mixture model produced an explaining-away effect that led to holistic processing (Figures 4E). This combination was key to simultaneous explanation of two important neural properties: tuning to a small number of facial features and face selectivity. That is, although the former property could be explained by sparse coding alone (Figure 10), the latter could not (Figure 4B) presumably since facial parts could accidentally be similar to object parts. However, when the sparse coding submodels for faces and objects were combined in the mixture model, the individual face units could be activated only if the whole input was interpreted as a face. In this sense, our theory interprets the face selectivity property as a signature of holistic processing. (It should be noted that the face selectivity may not be considered an “emergent” property of the model in the same sense as the tuning properties, since some kind of enhanced selectivity might well be expected by the introduction of a mixture model.) We also linked our model with more classical experiments on holistic processing by reproducing the tuning properties for partial or inverted faces (Figure 8). However, we could not prove the necessity of the mixture computation in these cases since the results without mixture were still consistent, albeit more weakly, with the experimental data.

Having explained known response properties, we can draw a few testable predictions of unknown properties from our theory. First, since face selectivity depends on the computational progress of stimulus interpretation as a face or as an object, we can predict delayed suppression in responses of face-selective neurons to non-face stimuli. Second, since face selectivity depends on the failure of stimulus interpretation as an object, we can predict loss of selectivity of face-selective neurons after deactivation of the object-selective region by muscimol injection or cooling.

Among the reported properties of face neurons in the monkey IT cortex, preferences to extreme features (in particular, monotonic tuning curves) were considered as a surprising property [4] since they were rather different from more typical bell-like shapes such as orientation and frequency tunings. We showed that our model explained quite well such extremity preferences (Figure 7). It is intriguing why our model face units had such property. First, we would like to point out that the facial features discussed here are mostly related to positions of facial parts and such features can be relatively easily encoded by a linear function of an image. This is not the case, however, for orientations and frequencies since encoding these seem to require a much more complicated nonlinear function, perhaps naturally leading to units with bell-like tunings. Second, we could speculate that the extremity preferences may be really necessary due to the statistical structure of natural face images, irrelevant to any particular details of our model. Indeed, even when we perform a very basic statistical analysis of principal components of face images (so-called eigenfaces, e.g., [34]), they look like linear representations of certain facial features, maximal in one extreme and minimal in the other extreme. However, this seems to be a rather deep question and fully answering it is beyond the scope of this study.

The results shown here relied on all computational components in mixture of sparse coding models, including inference computation of each sparse coding submodel and suppressive operations using computed posterior probabilities. Since these computations seem to be difficult to implement only with simple feedforward processing in the biological visual system, a natural assumption would be some kind of recurrent computation possibly involving feedback processing. While quite a few biologically plausible implementations have been proposed for sparse coding inference, e.g., [31, 35], we prefer here not to speculate how the mixture computation might be implemented, in particular, whether class information as in the top layer in our model might be represented explicitly in some cortical area or implicitly as some kind of mutual inhibition circuit between the face-selective and the object-selective regions in IT.

Related to the previous point, it would also be interesting whether or not similar results could be reproduced by a deep (feedforward) neural network model [6–12]. Note that, although face-selective units, tuning properties to head orientation, or behavioral properties on holistic face processing (such as the face inversion effect) have been discovered in some models [11–13, 36], no tuning properties to facial features like here have been reported yet. We particularly wonder whether the face-selective units in such models represent facial parts, since such parts are sometimes impossible to recognize correctly without any surrounding context if the input image does not contain enough detail, e.g., Figure 1B. While it is mathematically true that such nonlinear context-dependent computation could also be arbitrarily well approximated by a feedforward model, whether this can be achieved by a network optimized for image classification needs to be investigated empirically. In any case, however, we think that top-down feedback processing as formulated in our model would be a simpler and biologically more natural way of performing such computation.

Since we trained each submodel of our mixture model separately by face or object images, our learning algorithm was supervised, implicitly using class labels (“face” or “object”). This choice was primarily for simplification in the sense of avoiding the generally complicated problem of unsupervised learning of a mixture model. We do not claim by any means that face and object representations in the IT cortex should be learned exactly in this way. Nonetheless, the existence of such teaching signals may not be a totally unreasonable assumption in the actual neural system. In particular, since faces can be detected by a rather simple operation [37, 38], some kind of innate mechanism would easily be imaginable. This may also be related to the well-known fact that infant monkeys and humans can recognize faces immediately after eye opening [39, 40].

Early work on sparse coding concentrated on explaining receptive field properties of V1 simple cells in terms of local statistics of natural images [14, 15], following Barlow's efficient coding hypothesis [41, 42]. The theory was subsequently extended to explain other properties of V1 complex cells [17–19] and V2 cells [20–22]. The present study continues this approach to investigate higher visual representations, though a novel finding here is that an additional mechanism, a mixture model, is necessary to explain the neural properties discussed here. On the other hand, in computer vision, sparse-coding-like models have also been used for feature representation learning. In particular, the classical study on ICA of face images [34] may be related to the construction of our face sparse coding submodel, although the previous study reported global facial features as the resulting basis set [34]. (Because of this, it was once argued that parts-based representations require the non-negativity constraint [23]. However, it seems that such completely global ICA features may have been due to some kind of overlearning and, indeed, local feature representations were obtained when we used enough data as in Figure 3; we also confirmed this in the case with raw images.) Another relevant formalism is mixture of ICA models [43]. Although the idea is somewhat similar to ours, their full rank assumption on the basis matrix and the lack of Gaussian noise (reconstruction error) terms make it inappropriate in our case because the strong dimension reduction was essential for ensuring the face selectivity (Figure 11).

Our model presented here is not meant to explain all the properties of face neurons. Indeed, the properties explained here are a part of known properties of face neurons in the middle patch, which is in turn a part of the face network in the monkey IT cortex [25, 44, 45]. In the middle patch, face neurons are also tuned to contrast polarities between facial parts [46]. In more anterior patches, face neurons are tuned to viewing angles in a mirror-symmetric manner or invariant to viewing angles but selective to identities [47]. Further, all these neurons are invariant to shift and size transformation as usual for IT neurons [47]. Explaining any of these properties seems to require a substantial extension of our current model and is thus left for future research. Finally, since most detailed and reliable experimental data on the IT cortex concerns face processing, we hope that the principles, such as presented here, found in face processing could serve to elucidate principles of general visual object processing.

## Methods

### Model details

Our hierarchical model began with a bank of Gabor filters. The filters had all combinations of 10 × 10 center locations (arranged in a square grid within 64 × 64 pixels), 8 orientations (at 22.5° interval), 3 frequencies (0.25, 0.17, and 0.13 cycles/picels), and 2 phases (0° and 90°). The Euclidean norm of each Gabor filter with frequency *f* was set to *f* ^1.15^ (following 1/*f* spectrum of natural images) and the Gaussian width and length were both set to 0.4/*f*.

### Data preprocessing

As a face image dataset, we used a version of Labeled Faces in Wild (LFW) [28] where face alignment was already performed using an algorithm called “deep funneling” [29]. By this alignment, faces had a more or less similar position, size, and (upright) posture across images. The dataset consisted of about 13,000 images in total. Each image was converted to gray scale, cropped to the central square region containing only the facial parts and hairs, and resized to 64 × 64 pixels. Since many images still contained some background, they were further passed to a disk-like filter, which retained the image region within 30 pixels from the center and gradually faded the region away from this circular area. Finally, the pixel values were standardized to zero mean and unit variance per image.

As an object image dataset, we used Caltech101 [24]. We removed four image categories containing human and animal face images (Faces, Faces easy, Cougar face, and Dalmetian). The objects within the images were already aligned. The dataset consisted of about 8,000 images in total. Like face images, each image was converted to gray scale, cropped to square, resized to 64 × 64 pixels, passed to the above mentioned disk-like filter, and standardized per image.

For each class, we reserved 1,000 images for selectivity test and used the rest for model training.

### Learning details

To train the mixture model, we first processed the images with the energy detectors and then subtracted, from each data **x**, the dimension along the mean 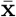 of all (face and object) data:

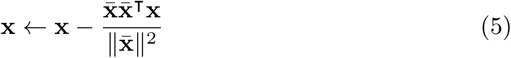

Although this operation was not quite essential, this had the effect of a linear form of contrast normalization suppressing a part of inputs with prominently strong signals; in fact, we observed that, without this operation, some elements of mean vectors **b***^k^* estimated as below became outrageously large.

Then, for each submodel for image class *k*, we learned the basis matrix **A***^k^* and the mean vector **b***^k^* in the following two steps:

1. perform strong dimension reduction using PCA [32] from 2400 to 100 dimensions while whitening;
2. apply overcomplete ICA [33] to estimate 400 components from 100 dimensions.

For overcomplete ICA, we used the score matching method for computational efficiency [33]. Formally, let **d***^k^* be the vector of top 100 eigenvalues (from PCA) sorted in descending order, **E***^k^* be the matrix of the corresponding (row) eigenvectors, and **R***^k^* be the weight matrix estimated by the overcomplete ICA. Then, using the filter matrix defined as

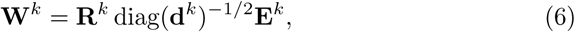

the basis matrix can be calculated as **A***^k^* = (**W***^k^*)^#^ (# is the pseudo inverse) and the mean vector as 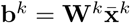 (where 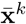 is the mean of all data of class *k*). Note that the signs of the filter vectors obtained from ICA are arbitrary; for the present purpose, we adjusted each sign so that all elements of **b***^k^* became non-negative.

### Theory of mixture of sparse coding models

A mixture of sparse coding models is similar to a classical mixture of Gaussians [30] in that it describes data coming from a fixed number of categories, but different in that each category is defined by a sparse coding model [14].

Formally, we assume an observed variable **x**: *R^D^*, a (discrete) hidden variable *k*: {1, 2,…, *K* }, and *K* hidden variables **y***^h^*: *R^M^* (*h* = 1, 2,…, *K*). Intuitively, **x** represents a (processed) input image, *k* represents the index of an image class (submodel), and **y***^h^* represents features (responses) for the class *h*.

We define the generative process of these variables as follows (see Figure 13 for the graphical diagram). First, an image class *k* is drawn from a pre-fixed prior *π_h_*: [0, 1] (where 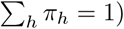:

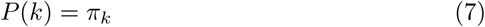

**Figure 13.**
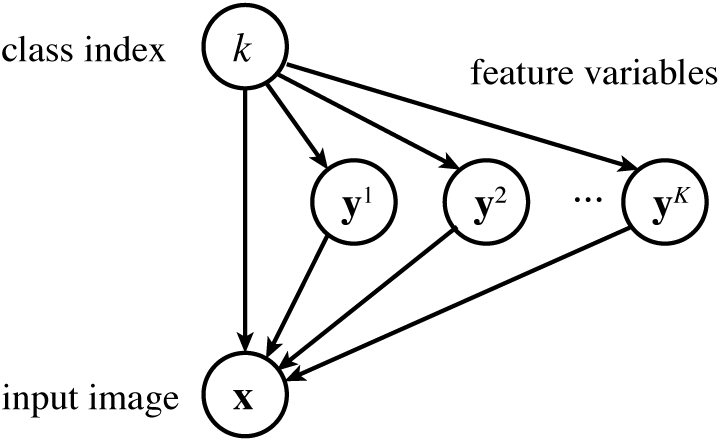
The graphical diagram for a mixture of sparse coding models. The variable *k* is first drawn from its prior, then each variable **y***^h^* is draw from a Laplace distribution depending on whether *h* = *k* or not, and finally the variable **x** is generated from a Gaussian distribution depending on **y***^k^*. (Note that, until *k* is determined, **x** is dependent on *k* and all of **y**^1^, **y**^2^, …, **y***^K^*.)

We call *k* here the generating class. Next, features **y***^k^* for the class *k* are drawn from the Laplace distribution with mean vector **b***^k^*: *R^M^* and a pre-fixed standard deviation λ (common for all dimensions)

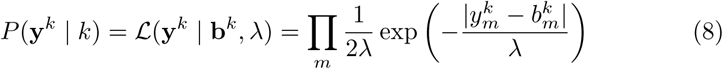

and an observed image **x** is generated from the features **y***^k^* by transforming it by the basis matrix **A***^k^*: *R^D × M^*, with a Gaussian noise of a pre-fixed variance σ^2^ added:

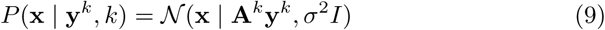

Here, **A***^k^* and **b***^k^* are model parameters estimated from data (see the section on Learning details above). Features **y***^h^* for each non-generating class *h* ≠ *k* are drawn from the zero-mean Laplacian

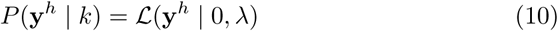

and never used for generating **x**. Altogether, the model distribution is rewritten as follows:

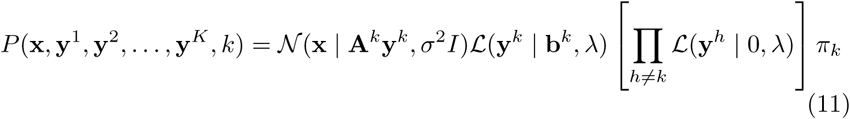

Since data are generated from the mixture of *K* distributions each of which is a combination of a Laplacian and a Gaussian similar to the classical sparse coding model [31], we call the above framework mixture of sparse coding models.

However, we depart from standard formulation of mixture models or sparse coding in two ways, motivated for modeling face neurons. First, since the feature variable **y***^h^* for the non-generating classes *h* ≠ *k* are unused for generating **x**, a standard formulation would simply drop the factor (10), leaving **y***^h^* unconstrained. However, our goal here is to model the responses of all (face or object) neurons for all stimuli (faces or objects). In fact, actual face neurons are normally strongly activated by face stimuli, but are deactivated by non-face stimuli, which is why our model uses a zero mean for non-generating feature variables. Second, the classical sparse coding uses a zero-mean prior [31], which is suitable for natural image patch inputs since their mean is zero (blank image) and this evokes no response like V1 neurons. However, the mean of face images is not zero and such mean face image usually elicits non-zero responses of actual face neurons. Therefore our model uses a prior with potentially non-zero mean **b***^k^* on the feature variable **y***^k^* for the generating class.

Given an input **x**, how do we infer the hidden variables **y***^h^*? Since evoked response values of neurons that are experimentally reported are usually the firing rates averaged over trials, we model these quantities as posterior expectations of the hidden variables. Since exact computation of those values would be too slow, we use the following approximation (see the derivation in the section on Approximating posterior later).

1. For each image class *k*, compute the MAP (maximum a posteriori) estimates of the feature variables **y**^1^, **y**^2^,…, **y***^K^*, conditioned on the class *k*:

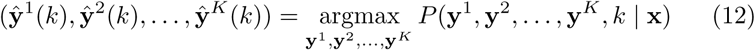
2. Compute the approximate posterior probability of each image class *k*:

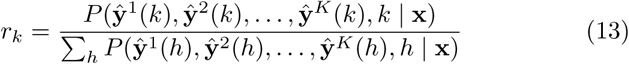
3. Compute the approximate posterior expectation of each feature variable *k*:

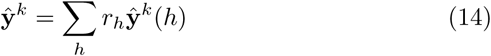

Note that, in equation (12), the feature variables for non-selected classes are always exactly zero:

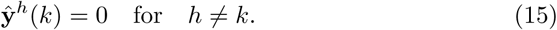

Therefore, even though an alternative approach would be to model neural responses by the MAP estimates of feature variable for the best image class, this may be too radical since responses becoming absolutely zero are a little unnatural.

The Bayesian inference described in the section on “Model” can be derived from steps 1 to 3 above in a straightforward manner using the model definition (11) and the property (15).

### Approximating posterior

Given an input *x*, we intend to compute the posterior expectations of each **y***^h^*:

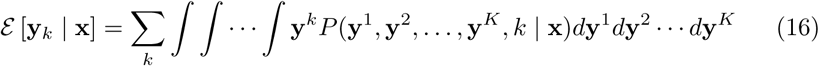

Direct computation of this value is not easy. Note, however, that, from the definition of the model (equation 11), the posterior distribution has a single strong peak for each class *k*, with variances more or less similar across all classes. Therefore we approximate the posterior probability by

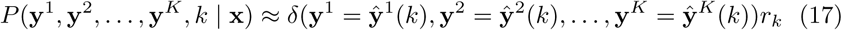

where 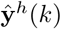 is the MAP estimate of **y***^h^* when the selected image class is *k* (equation 12) and *r_k_* is the relative peak posterior probability for the class *k* (equation 13). Here, δ(·) is the delta function that takes infinity for the specified input value and zero for other values. Substituting the approximation (17) into equation (16) yields equation (14).

### Simulation details

Cartoon face images were created by using the method described by Freiwald et al. [4]. Each face image was drawn as a linear combination of 7 facial parts (outline, hair, eye pair, iris pair, eyebrows, nose, and mouth). The facial parts were controlled by 19 feature parameters: (1) face aspect ratio (round to long), (2) face direction (left to right), (3) feature assembly height (up to down), (4) hair length (short to long), (5) hair thickness (thin to thick), (6) eyebrow slant (angry to worried), (7) eyebrow width (short to long), (8) eyebrow height (up to down), (9) inter-eye distance (narrow to wide), (10) eye eccentricity (long to round), (11) eye size (small to large), (12) iris size (small to large), (13) gaze direction (11 *x*-*y* positions), (14) nose base (narrow to wide), (15) nose altitude (short to long), (16) mouth-nose distance (short to long), (17) mouth size (narrow to wide), (18) mouth top (smily to frowny), and (19) mouth bottom (closed to open). Note that the first three parameters globally affected the actual geometry of all the facial parts, while the rest locally determined only the relevant facial part. See Figure S3 for example images.

Following the method in the same study [4], we estimated three kinds of tuning curves: (1) full variation, (2) single variation, and (3) partial face. For full variation, a set of 5000 cartoon face images were generated while the 19 parameters were randomly varied. For each unit and each feature parameter, a tuning curve at each feature value was estimated as the average of the unit responses to the cartoon face images for which the feature parameter took that value. The tuning curve was then smoothed by a Gaussian kernel with unit variance. To determine the significance of each tuning curve, 5000 surrogate tuning curves were generated by destroying the correspondences between the stimuli and the responses. Then, a tuning curve was regarded significant if (1) its maximum was at least 25% greater than its minimum and (2) its heterogeneity exceeded 99.9% of those of the surrogates, where the heterogeneity of a tuning curve was defined as the negative entropy when the values in the curve were taken as relative probabilities.

For single variation, a tuning curve for a feature parameter at each value was estimated as the response to a cartoon face image for which the feature parameter took that value and the other were fixed to standard values. The standard parameter values were obtained by a manual adjustment with the stimuli used in the experiment [4, Suppl. Fig. 1]. For partial face, cartoon face images with only one facial part (hair, outline, eyebrows, eyes, nose, mouth, or irises) were created. Each tuning curve for each feature parameter was obtained similarly to single variation, except that only the relevant facial part was present in the stimulus.

## Supporting information

**Figure S1.**
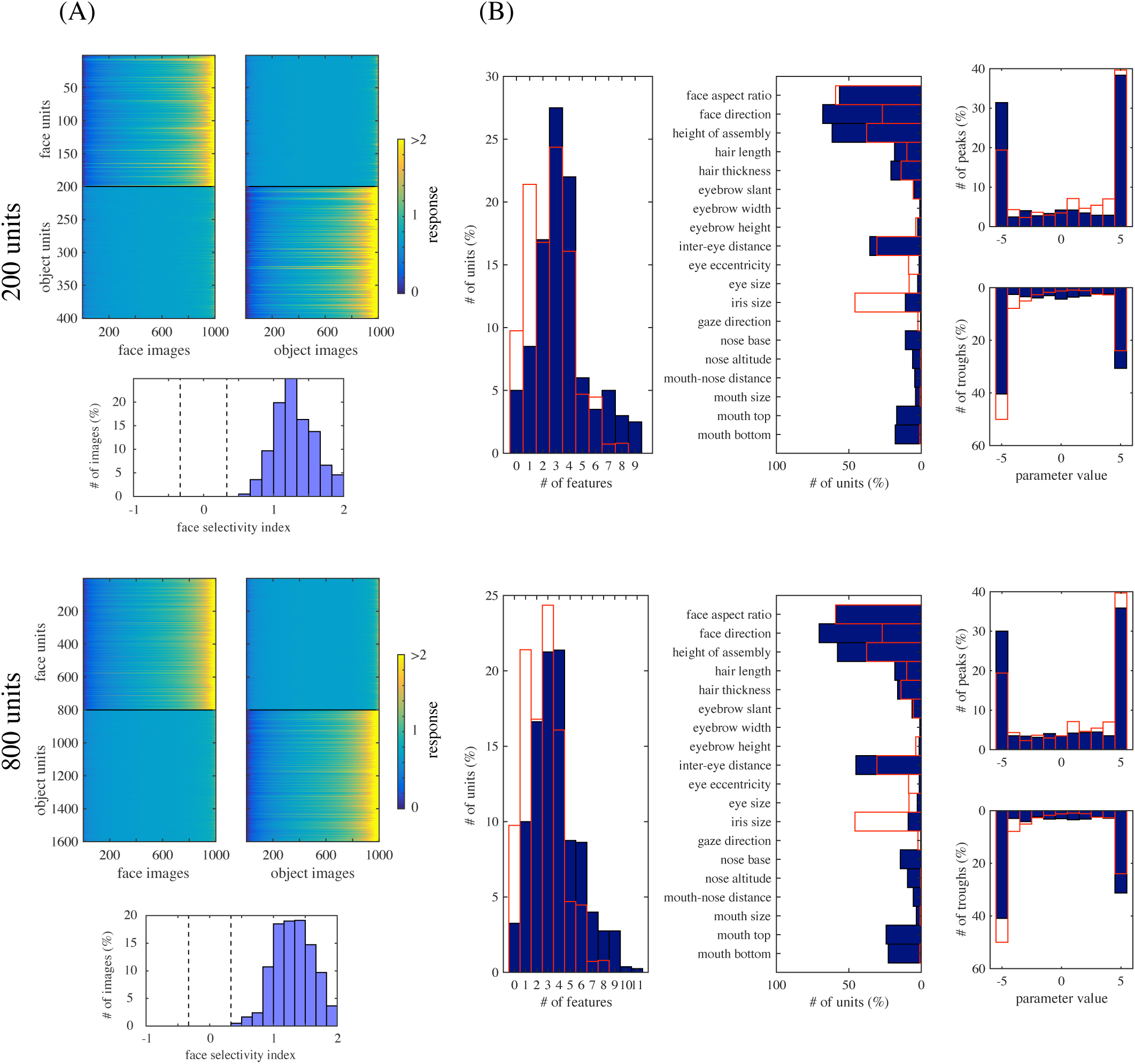
Control simulations varying the number of units. A mixture model was constructed in the same way as the original one, except that each submodel here had 200 units (upper half) or 800 units (lower half). (A) The responses of model face units and object units to natural face images (left) or natural object images (right), together with the distribution of face-selective indices for the face units (bottom); compare these with Figure 4A and D (blue). (B) The distributions of the numbers of significantly tuned features (of cartoon faces) per unit (left), of numbers of significantly tuned units for each feature parameter (middle), of peak and trough parameter values (right); compare these with Figures 6 and 7B. Overlaid red boxes are replots of corresponding experimental data [4].

**Figure S2.**
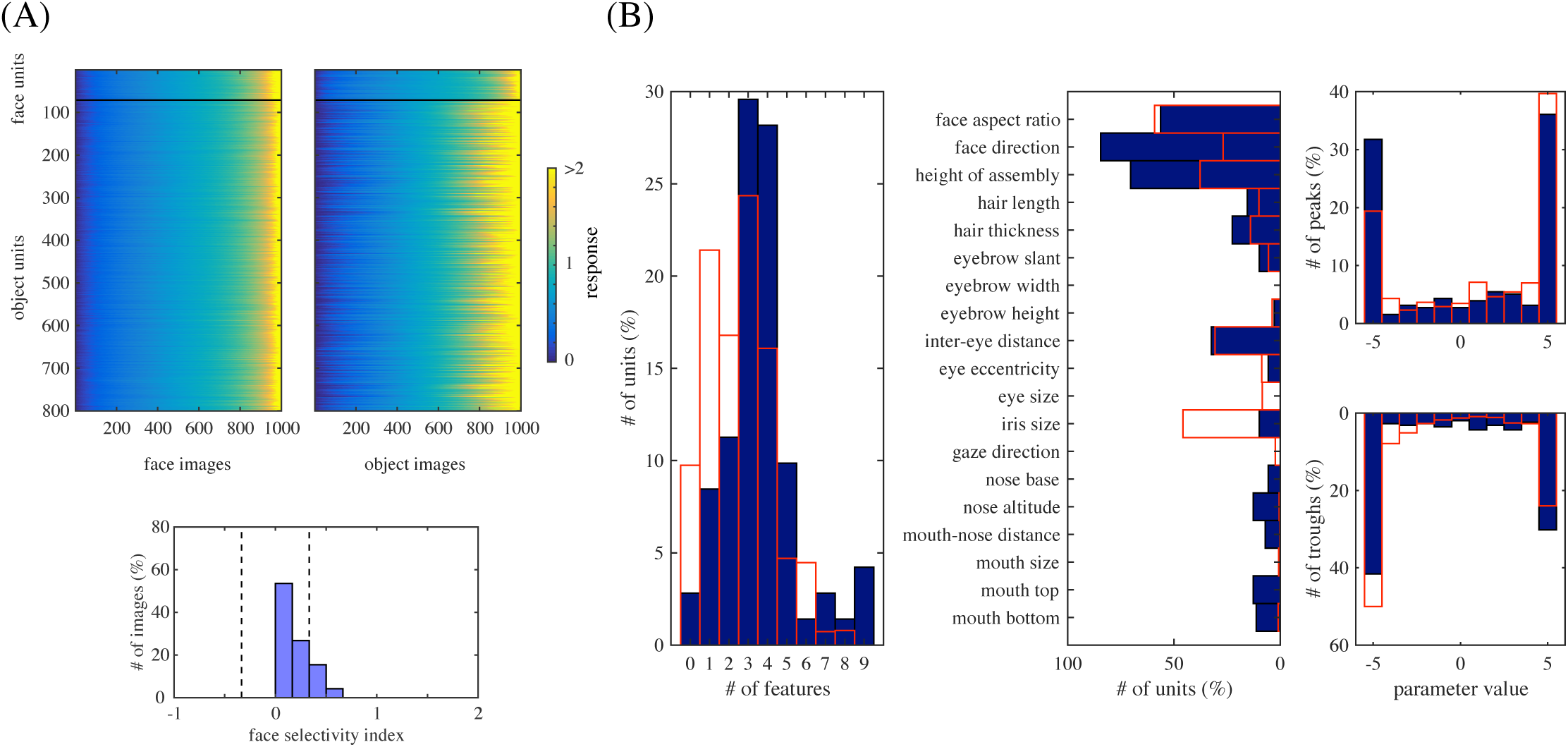
Control simulation with a single sparse coding model. A single sparse coding model with 800 units was constructed on top of the same energy model and trained with an ensemble of face and non-face images. In the resulting model, only 71 units gave larger average responses to face images than non-face images. The response properties of these units are shown. (A) The responses of face and object units to face images (left) or object images (right), with the distribution of face-selective indices for the face units (bottom). No prominent selectivity like in Figure 4A can be observed; the result is more similar to Figure 4B. (B) The distributions of the numbers of significantly tuned features per unit (left), of numbers of significantly tuned units for each cartoon face feature parameter (middle), of peak and trough parameter values (right); compare these with Figures 6 and 7B as well as Figure 10 (cyan curves). Overlaid red boxes are replots of corresponding experimental data [4].

**Figure S3.**
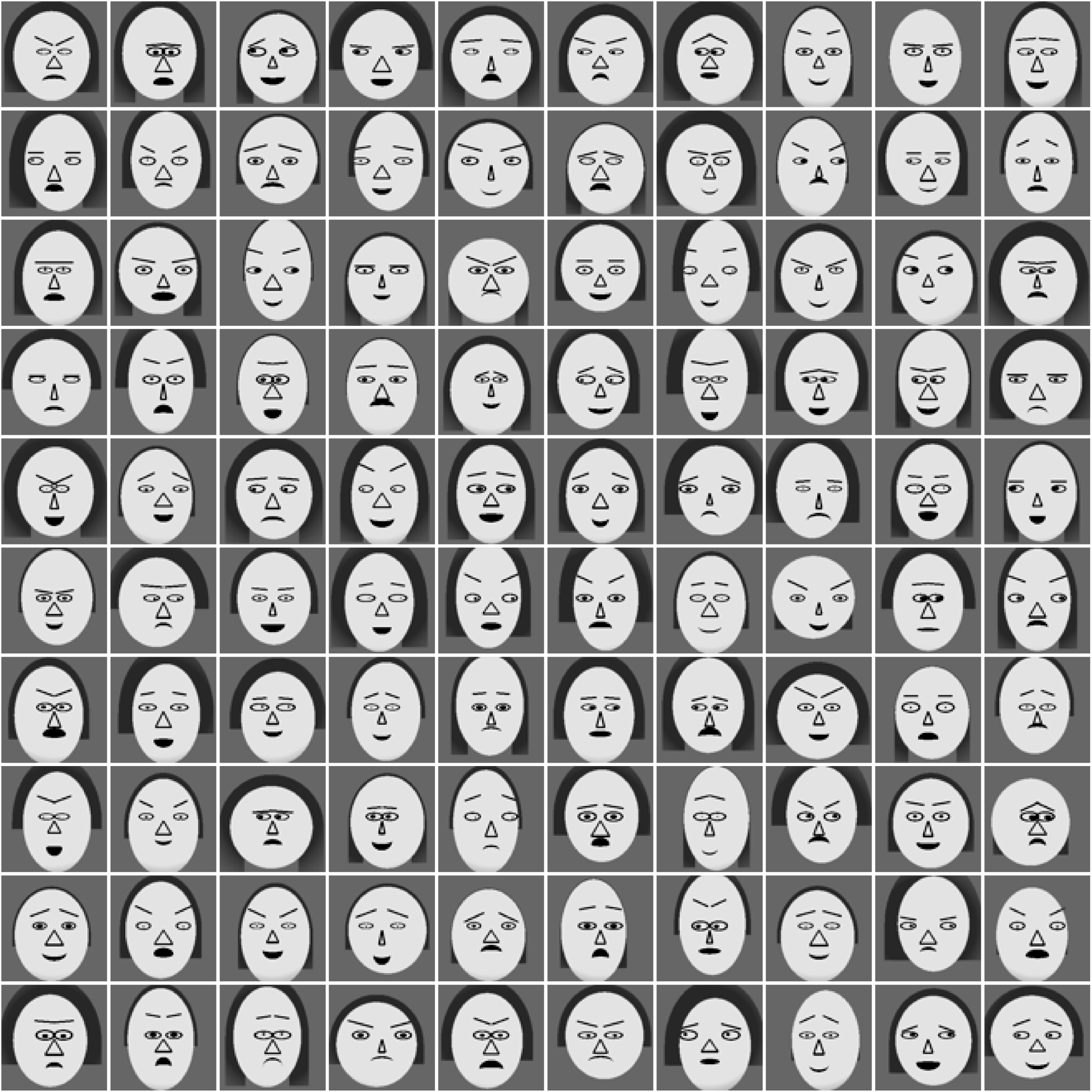
Random examples of cartoon face images.

